# Androgen Receptor driven gene score identifies tumors with Epithelial to Mesenchymal Transition features in triple negative breast cancer

**DOI:** 10.1101/2023.02.21.529333

**Authors:** Savitha Rajarajan, VP Snijesh, CE Anupama, Madhumathy G Nair, D Apoorva, M Chandrakala, Sharada Patil, Vidya P Nimbalkar, Annie Alexander, Maalavika Pillai, Mohit Kumar Jolly, Radhakrishnan Sabarinathan, Rakesh S Ramesh, BS Srinath, Jyothi S Prabhu

## Abstract

**Background:** Androgen receptor (AR) is considered marker associated with better prognosis within hormone receptor positive tumors. Its role in triple negative tumors however is controversial showing both better and worse prognosis and different methods are used for identification of AR driven tumors. Conflicting results could be due to intrinsic molecular differences or scoring method for AR positivity. We attempted to develop an AR driven gene score and examined its utility in subtypes of breast cancer (BC).

**Methods:** A bioinformatic pipeline was constructed and applied on publicly available microarray data sets obtained from AR positive BC cell lines treated with dihydrotestosterone (DHT). Expression levels of genes identified through the pipeline along with set of epithelial mesenchymal transition (EMT) markers, proliferation associated genes and enzymes involved in intracrinal androgen metabolism was evaluated in a cohort of Indian Breast cancer patients. Tumors were divided into AR high and low based on the gene score and association with clinical parameters, circulating androgens, disease free survival, proliferation and EMT markers were examined, all results were further validated in external public datasets.

**Results:** AR driven gene score was calculated as average expression of the 6 genes selected through bioinformatic analysis. 53% (133/249) tumors were classified as AR high by the gene score and had significantly better clinical parameters such as higher age, smaller tumor size, lower grade and lower proliferation. Tumors with high AR driven gene score had significantly better disease-free survival (mean survival time of 86.13 vs 72.69 months, log rank p=0.032) when compared to the AR low tumors. A subset analysis within TNBC (N=66) showed 36% (24/66) were AR high and had significantly higher expression of EMT markers (p=0.024). Though circulating levels of total testosterone was not different between the groups, intratumoral levels of 5 alfa reductase (SRD5A1) was significantly high in tumors with high AR driven score.

**Conclusion:** Role of AR in breast cancer is debatable and difficult to decipher with protein detection alone. Our results support the context dependent function of AR in driving better prognosis, while identifying its role in driving EMT within TNBC tumors.

## Introduction

Breast cancer (BC) is the most common tumor in the female population and highest in terms of incidence and mortality in Indian women(1). It is characterized by heterogeneity both at molecular and clinical level. The main breast cancer subtypes have been classified according to their molecular profile with identification of targets for treatment which have significantly improved the prognosis for hormone receptor positive (HR+HER2-) and HER2 amplified (HER2+) tumors(2). Triple negative breast cancers (TNBC) on the other hand are mainly treated with cytotoxic chemotherapy due to lack of therapeutic targets. There is a need to identify alternate therapeutic targets to diminish drug resistance and improve clinical benefits for this subtype of BC.

Androgen receptor (AR) plays an important role in the biology of breast cancer and is considered a useful marker for prognosis. It is implicated in BC development and its function vary across different subtypes of BC. Though frequently expressed in all subtypes of BC, the expression of AR and its pathways differ within subclasses. Evidence from published literature supports its role as tumor suppressor within HR+HER2-BC and its presence is associated with better prognosis within this subtype(3). Its role in TNBC has been debatable and shows both better and worse prognosis based on the ethnic population and different methods used for identifying the AR driven tumor(4–8). Prognostic role of AR within TNBC is uncertain due to use of different antibodies and cut off values used for protein assessment(9,10). A recent multi-institutional study(11) shows AR protein status by IHC alone is not a reliable marker for prognosis and variations in the expression levels of AR protein was observed across populations despite using well standardized uniform methods. Conflicting results could also be due to intrinsic molecular differences within AR based signalling within BC. Role of AR and androgens in carcinogenesis and promoting metastasis through epithelial to mesenchymal transition (EMT) is well established in prostate cancer(12).Similar trends of AR in promotion of EMT has been recently shown in *in vitro* models of BC(13–15). Moreover, AR is also considered a drug target in BC and AR targeted therapies have induced varied tumor responses in clinical trials(16–19). Therefore, deeper examination of the pathways regulated by AR is necessary to get an in-depth understanding of the molecular mechanisms involved in AR mediated signalling in breast cancer.

Gene expression profiles have shown evidence in identifying disease subtypes and have been more useful than employing single markers and hence are widely used for clinical application(20–22). Using multiple markers to derive AR regulated gene signatures could help identify breast tumors truly driven by AR and are amenable to anti androgen therapies. Here, we attempted to develop an AR driven gene score, examined its association with the clinicopathological and luminal features and its role in the commonly altered pathways of proliferation and EMT within BC.

## Material and Methods

### Bioinformatic methods to identify the AR driven genes

#### Collection of datasets

The gene expression profiles of AR positive breast cancer cell lines with series identifier GSE61368(23) and GSE28305(24) were retrieved from Gene Expression Omnibus (GEO) database. GSE61368 was derived from ZR751 cell line (ER+/AR+) cotreated with dihydrotestosterone (DHT) and estradiol for 16h at 10nM concentration.GSE28305 was derived from MDA-MB-453 cell line (ER-/AR+) treated with 10nM DHT for 16h. The detailed sample information is given in the Supplementary Table S1. The curated genes from various studies related to the androgen or estrogen in breast cancer were collected and termed as ‘base genes.

#### Data analysis and prioritization of candidate genes

Analysis on the expression data was performed using R package *limma*(25). To standardize and reduce the technical noise in the probe level data, the raw signal values of each probe sets were normalized using Robust Multiarray Average (RMA)(26) algorithm. The differentially expressed genes (DEGs) between the control and treatment groups were filtered based on the p-value (p≤0.05) significance.

#### Protein Interaction Map and Network Analysis

The list of DEGs and ‘base genes’ were mapped to the Human Integrated Protein-Protein Interaction Reference (HIPPIE) database(27,28) for constructing Protein Interaction Map (PIM). All the protein interactions of DEGs and ‘base genes’ obtained from the gene expression analysis were extracted with an association score of ≥ 0.4 to create PIM. Visualization and calculation of topological parameters of PIM were performed using Cytoscape (version 3.8.2)(29). We adopted an approach, which has been formerly applied by Rakshit et al., 2014(30) to identify the hubs. The degree centrality (DC) cut-off threshold formula for choosing the hub protein is defined as:

Hubs = M + (2 x SD), where M=Mean Degree across the genes and SD = Standard deviation of the degree across genes.

From the PIM, genes and their primary partners that belong to hubs and base genes were extracted to decompose the complex interactome PIM to a Significant Protein Interaction Map (sPIM).

#### Calculation of Semantic Similarity

Using encoded evidence in the gene ontology (GO) hierarchy, the functional similarity between the gene pairs in the subnetwork sPIM was assessed using R package *GoSemSim(31*). In this study, we used Wang’s similarity metric (32)to compare the biological process (BP) hierarchy. Next, we filtered the gene pairs in which one among the gene in the pair has at least absolute fold change of 1.4. To retain genes which have differential expression and were removed due to lack of connectivity, absolute fold change with specific threshold was screened from the initial list of significant genes and those which fall above top 75 quantile were screened and included in the final list.

### Cohort Details

Tumor samples were chosen from a retrospective cohort of 244 women with primary breast cancer including 5 women with bilateral tumors. These samples were collected as part of an observational longitudinal study from two tertiary cancer care hospitals in Bangalore, India between 2008 and 2013 and these women were followed-up for up to 9 years, with a total loss to follow up of less than 5%, and a median follow-up duration of more than 72 months. Informed consent was obtained from all the patients to use their tissue and blood sample for research and the study was approved by the ethical committee of both institutions. Information on clinical variables like age, grade, tumor size, lymph node status, stage of the disease with ER, Progesterone receptor (PgR) and HER2 were obtained from their clinical records (Supplementary Table S2). Formalin fixed paraffin embedded (FFPE) blocks from tumor tissue having more than 50% of the area of representative tumor were selected for the study.

### Immunohistochemistry of AR

Immunohistochemistry for AR was done on each of the tumor sections as per standard protocol using the Ventana BenchmarkXT staining system (Ventana Medical Systems, Tucson, AZ, USA), the detailed methodology is described in our previous publication(33), Primary antibody for AR(Clone AR 441, DAKO, dilution at 1:75) was used with positive and negative controls run for each batch. Two pathologists scored the staining for AR protein independently and arrived at a final score. Nuclear staining in more ≥1% of tumor cells was considered as positive.

We also accessed tissue micro array sections of an independent cohort (N=107) and examined the presence of AR and ZEB1 using the primary antibody for ZEB1 (Clone E2G6Y, Rabbit mAb, CST, Cat #70512) at a dilution of 1 in 200. Any staining in ≥1% of tumor cells or tumor associated fibroblasts was taken as positive for ZEB1.

### Estimation of testosterone

The estimation of total testosterone in serum samples of the selected breast cancer patients was done by a chemiluminescence based immunoassay method using the Abbott Architect ci8200 (Integrated) & i2000 (Immunoassay) instrument, detailed methodology described in our previous publication (33). The serum samples were collected prior to surgery or following surgery from 154 breast cancer patients.

### Gene expression by real time PCR

Total RNA extraction was done using the Tri Reagent protocol according to manufacturer’s instructions (Sigma Aldrich # T9424) from two 20 μm sections from the selected tumor block following the methods published previously. 500ng of total RNA was reverse transcribed to cDNA using high-capacity cDNA conversion kit from Thermofisher scientific (Cat # 4322171) as per manufacturer’s instruction.

Primers were designed for the AR driven genes (CYP4Z1, TFAP2B, ABCC11, SOCS2, GADD45G, ZNF689,ID1,PIP,UGT2B11,KCNMA1,SEC14L2 and DOCK2), proliferation related genes (BIRC5, ANLN CENPF and UBE2C), EMT related genes (SNAI2,TWIST1,ZEB1 and ZEB2) and genes coding for enzymes involved in androgen and oestrogen synthesis (SRD5A1 and CYP19A1) using primer 3 plus software and further validated on ensemble genome browser, NCBI blast and UCSC genome browser. The primers were synthesized by Juniper Life sciences, Bangalore, India. The details of the primer sequences are given in the supplementary table S3. The methods used for nucleic acid extraction, quantitative PCR (qPCR)and selection of housekeeping genes (HKG) and the quality control criteria for inclusion of samples in the analysis is described in detail in previous publication(34). Relative normalised expression was calculated for each gene as published in our previous methods(35)

To derive a proliferation score, a logistic regression model was constructed using proliferation related genes namely, BIRC5, ANLN CENPF, UBE2C and Ki67 protein as determinant. EMT score was taken as the average expression of EMT related genes. We also accessed the ER probability score for these tumors in the cohort derived according to methods described previously(36).

### External data sets accessed

Gene expression data (cDNA microarray profiling, Illumina HT-12 v3 platform) from Molecular Taxonomy of BRCA International Consortium (METABRIC) project was retrieved from the cBioPortal (37) (www.cbioportal.org/).

### Statistical methods

Descriptive analysis was done to evaluate the cohort characteristics and those between high and low AR score groups. Difference in the clinical variables between high and low AR groups was tested by independent Student’s t-test or Mann Whitney U test for continuous variables and chi-square test was done for categorical variables. Concordance between the AR driven score and protein was estimated by receiver operating characteristic (ROC) curve analysis. Kaplan-Meier survival curves and log rank tests were used to compare survival between the high and low AR score groups. Disease free survival (DFS) was calculated as the time from the date of first diagnosis to the time when a local or distant recurrence occurred. Patients with no event or had death due to non-breast cancer related causes were right censored. All tests were two tailed and P-value <0.05 was considered statistically significant. All statistical analyses were done on statistical software XLSTAT version 2022.1.2 and R software version 3.6.3.

## Results

### Deriving the AR driven gene score

We derived the AR driven genes through bioinformatic pipeline as described in the methods from publicly available microarray data sets obtained from ER+AR+ BC cell line (ZR-75-1) and ER-AR+ (MDAMB-453) BC cell line, both treated with dihydrotestosterone. Genes identified through the method were classified based on the context of ER expression (in presence of ER, independent of ER and absence of ER) as shown in Figure 1. Pathway analysis showed commonly regulated genes between both cell lines (ER+AR+ and ER-AR+), identified pathways related to androgen receptor and androgens, while gene sets derived from only ER-AR+ cell line detected pathways unrelated AR or androgens. Therefore, gene sets derived from presence of ER (35 genes) and independent of ER context (19 genes) were taken ahead for further analysis. A total of 54 genes were chosen and absolute fold change cut-off of 2 was used to arrive at the final set 12 genes. We further evaluated the transcript abundance of the chosen set of 12 genes (gene list in the supplementary table 3)

**FIGURE1:**
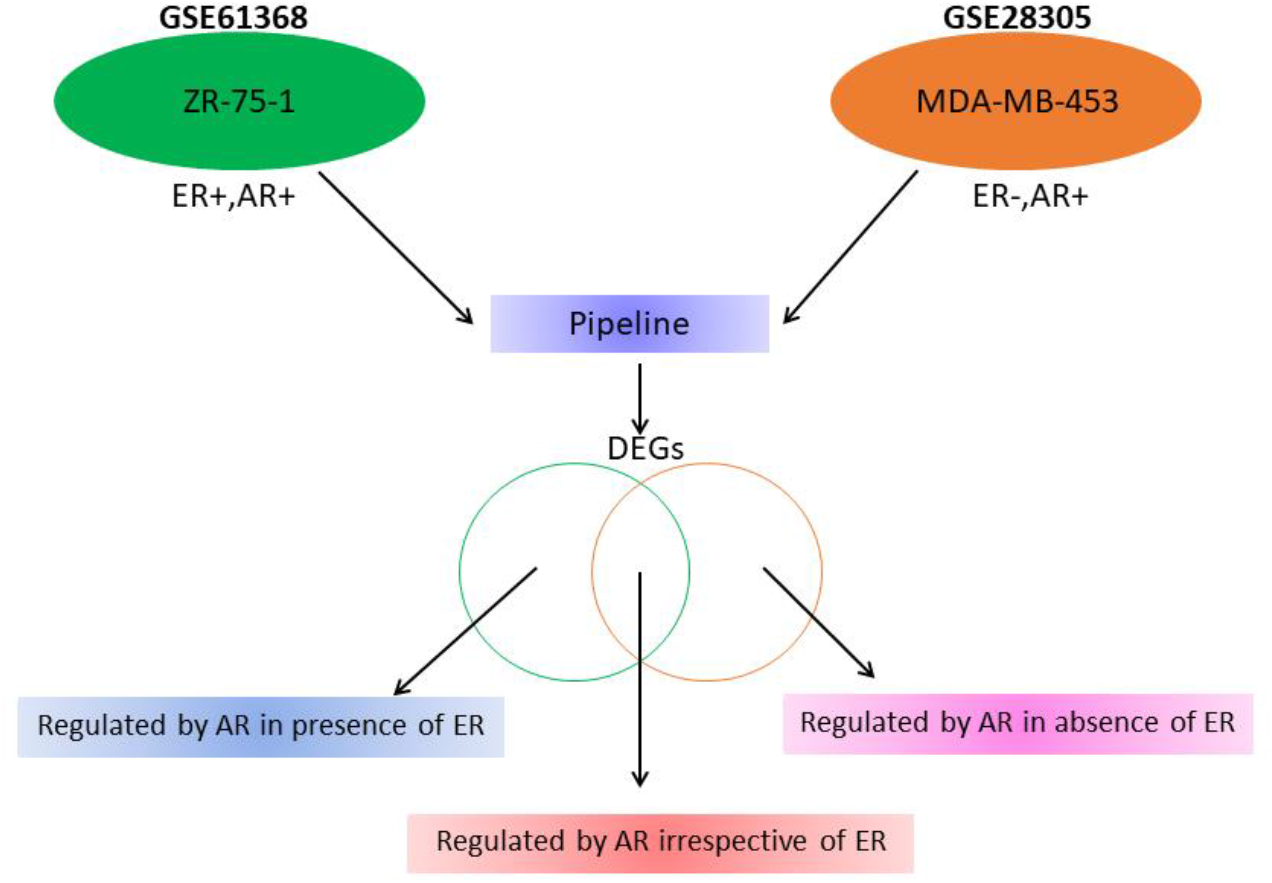
Bioinformatic workflow to arrive at the three sets of AR driven genes under different context of ER

Only 6 of the 12 genes (CYP4Z1, TFAP2B, ABCC11, PIP, KCNMA1 and SEC14L2) had higher fold change and positive correlation with AR transcript. AR driven gene score was calculated as average expression of these genes and this score had significant concordance with AR protein by ROC analysis (AUC-0.65, p=0.001).

### Prognostic value of AR driven gene score

The AR driven gene score ranged from 4.66 to 17.62 with a mean value of 10.51 and median of 10.6. The mean cut-off was taken to divide the tumors into AR high and AR low. 53% (133/249) tumors were classified as AR high by the gene score and had significantly better clinical parameters such as higher age, smaller tumor size, lower grade, higher proportion were post-menopausal (p<0.05), lower proliferation score (p=0.016) and higher ER probability score (p=0.007) as shown in Table 1.

**Table 1:**
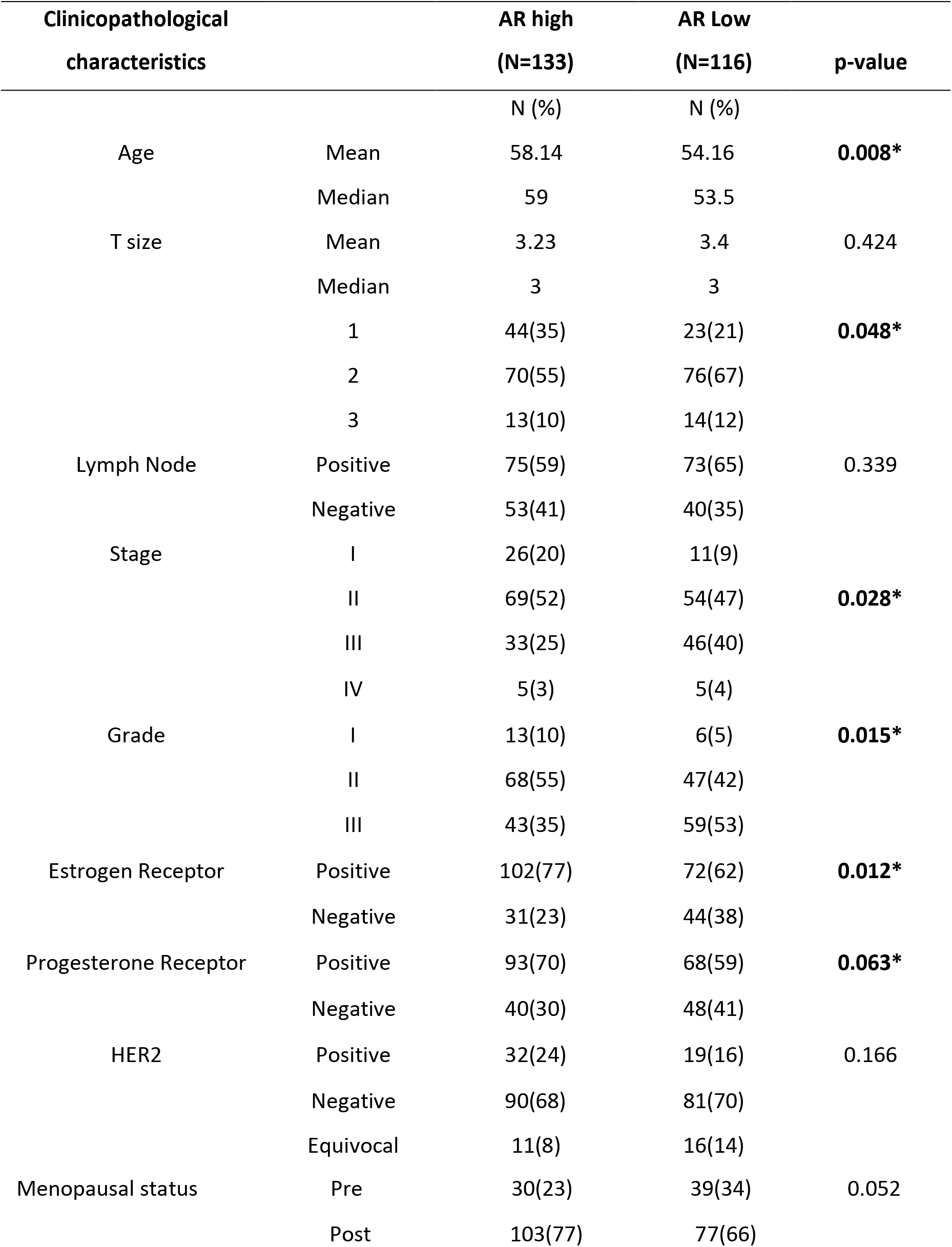
Comparison of clinical variables between high and low AR groups in all tumors(N=249)

AR protein status from IHC was available for 165/249 tumors. Of the 165 tumors, only 60 tumors were positive for AR protein (36%). As reported earlier(33), Kaplan Meier survival analysis between AR positive and negative tumors showed no significant difference in survival between the two groups, indicating that AR protein alone is not a good prognostic marker in our cohort. Disease free survival in the tumors with high AR driven gene score however was significantly better (mean survival time of 86.13 vs 72.69 months, log rank p=0.032) when compared to the AR low tumors, clearly demonstrating it to be a better prognostic indicator than the AR protein (Figure 2). A subset analysis of disease-free survival within the subtypes such as Hormone receptor positive tumors and Triple negative (TNBC) tumors however was not different.

**FIGURE 2:**
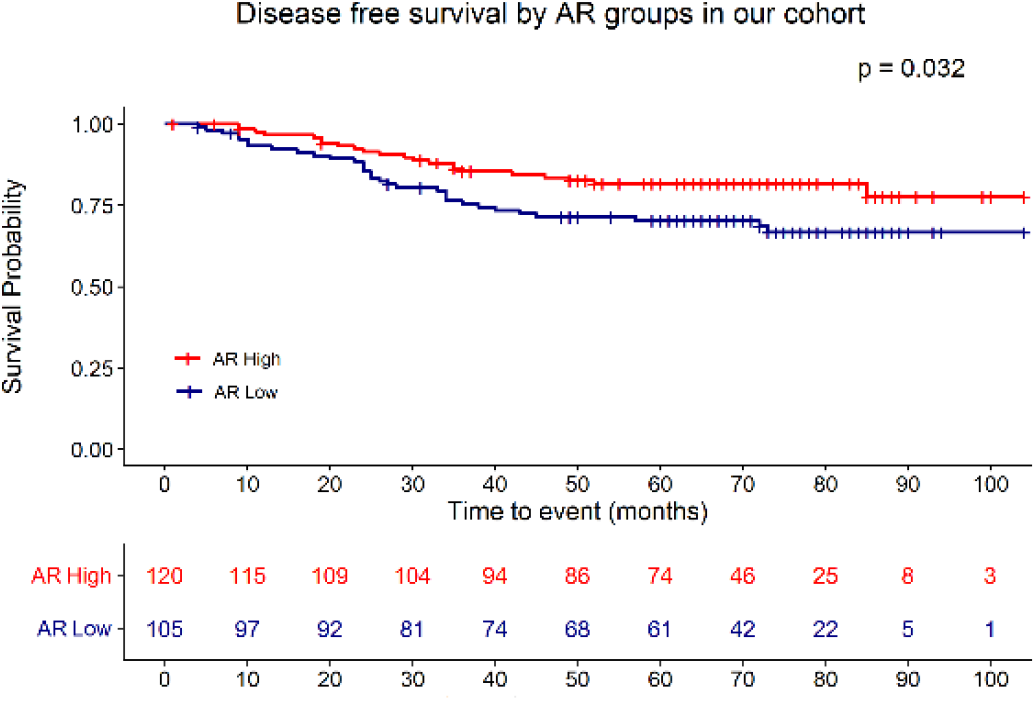
Kaplan Meier survival analysis showing the disease free survival between AR high and AR low groups in all tumors in our cohort.

### AR driven TNBC have high expression of SRD5A1

Intracrinal levels of the steroid hormones are known to influence the signalling through steroid receptors than their circulating levels. Therefore, we examined the expression levels of androgen synthesizing enzyme, SRD5A1 and estrogen synthesizing enzyme CYP19A1 within all tumors. SRD5A1 catalyses the conversion of testosterone to DHT and CYP19A1 catalyses the conversion of testosterone to estradiol. We observed AR driven tumors had a significant higher expression of SRD5A1 in all tumors and within the TNBC subtype (p=0.03 and p=0.04) (Figure 3A and 3B). No changes were seen in the level of SRD5A1 expression in HR+HER2- or HER2+ tumors. No significant difference was observed in the CYP19A1 levels between the AR high and low tumors. Though the SRD5A1 level was significantly different, no difference was observed in the circulating levels of total testosterone between the AR high and low group of tumors.

**FIGURE 3:**
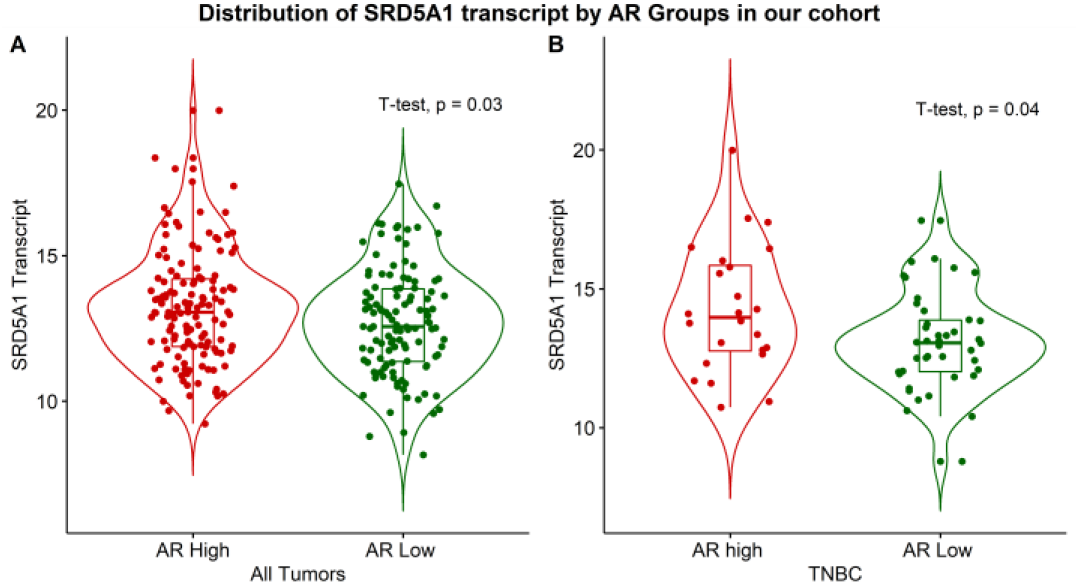
SRD5A1 levels in AR high and low groups in our cohort. (A) Distribution of SRD5A1 transcript in all tumors (B) Distribution of SRD5A1 transcript in TNBC tumors

### Tumors with high AR driven gene score have higher expression of EMT markers

AR is shown to induce EMT within prostate cancer while its association with EMT in BC is not known. We examined the distribution of EMT score (derived as mean expression of chosen key EMT genes namely, TWIST1, ZEB1, ZEB2 and SNAI2) between the high and low AR driven tumors and observed tumors with high AR driven gene score had significant higher EMT score than the low AR tumors (p=0.017) as seen in Figure 4A.

**FIGURE 4:**
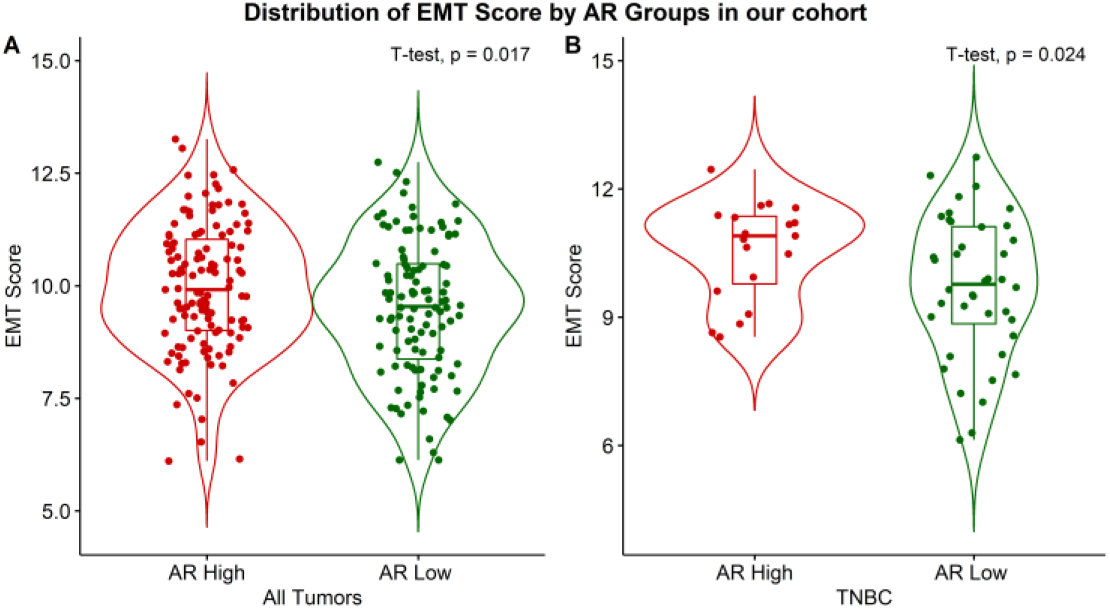
EMT score in AR high and low groups in our cohort. (A) Distribution of EMT score in all tumors (B) Distribution of EMT score in TNBC tumors

Next, we wanted to examine if the influence of AR on EMT is confined within any subtypes of breast cancer and hence divided the tumors into HR+HER2-(N=132), HER2+(N=51) and TNBC(N=66). Subset analysis within HR+HER2- and HER2+ tumors did not show any significant association between the two scores and no difference was observed in the EMT score between the AR high or low tumors.

In the TNBC tumors(N=66), 36%(24/66) had high AR driven score and these tumors showed favorable features such as higher age, smaller tumor size and significant lower proliferation score (p=0.008) and higher ER probability score (p<0.0001) (Table 2) suggestive of luminal associated features. However, tumors with high AR driven score had significant higher EMT score (p=0.024) than the AR low tumors (Figure 4B) indicating association of AR with EMT is confined to TNBC tumors alone.

**Table 2:** Comparison of clinical variables between high and low AR groups in TNBC tumors (N=66)

| Clinicopathological characteristics |  | AR high<br>(N=24) | AR Low<br>(N=42) | p-value |
| --- | --- | --- | --- | --- |
|  |  | N (%) | N (%) |  |
| Age | Mean | 55.33 | 50.97 | 0.143 |
|  | Median | 55 | 52 |  |
| T size | Mean | 3.45 | 3.6 | 0.685 |
|  | Median | 2.9 | 3.5 |  |
|  | 1 | 6(27) | 6(15) | 0.438 |
|  | 2 | 14(64) | 29(71) |  |
|  | 3 | 2(9) | 6(14) |  |
| Lymph Node | Positive | 12(52) | 20(49) | 0.794 |
|  | Negative | 11(48) | 21(51) |  |
| Stage | I | 5(21) | 5(12) | 0.439 |
|  | II | 10(42) | 24(57) |  |
|  | III | 7(29) | 12(29) |  |
|  | IV | 2(8) | 1(2) |  |
| Grade | I | 4(18) | 1(2) | 0.103 |
|  | II | 8(36) | 17(43) |  |
|  | III | 10(44) | 22(55) |  |
| Menopausal status | Pre | 7(29) | 15(36) | 0.585 |
|  | Post | 17(71) | 27(64) |  |

Crosstalk between major EMT regulating transcription factor ZEB1 and AR has been reported(38). The positive association of EMT score in AR high tumors urged us to examine the expression of ZEB1 in these tumors. A significant positive correlation was observed between ZEB1 transcript and AR driven gene score within all tumors (Pearson’s r=0.329, p<0.0010) and TNBC (Pearson’s r=0.409, p=0.001). We also examined the distribution of ZEB1 protein on the BC tumor cores in an independent cohort of tumors and found its expression predominantly in the stromal compartment of the tumor (Figure 5). 52/99 (53%) of tumors in the TMA were positive for AR. Comparison of the expression pattern of ZEB1 between AR positive and negative tumors showed higher expression of ZEB1 in AR positive tumors (p-0.065). Only 2/19(11%) of TNBC tumors were AR positive and we did not observe any difference in ZEB1 expression between AR positive and negative tumors within TNBC.

**FIGURE 5:**
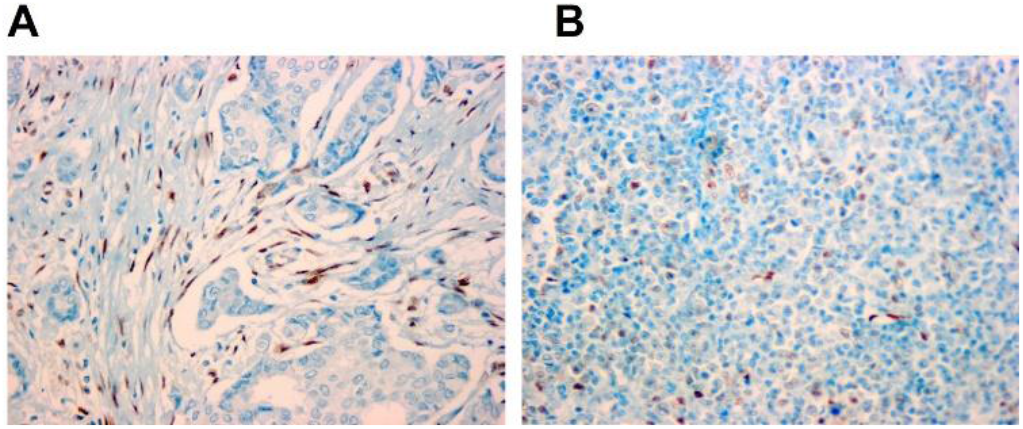
Representative IHC image of ZEB1 protein expression at 20X (A) In stroma and (B) In tumor

### Validation in external cohorts

To validate our findings, we accessed the METABRIC cohort with a total of 1904 tumors. Of these, 1369 tumors were HR+HER2-, 299 were TNBC and 236 were HER2+. As observed in our cohort, the 6 AR driven genes were significantly and positively correlated with AR transcript in the METABRIC cohort and the AR driven gene score was calculated by taking the average of 6 AR driven genes. This score ranged from 5.37 to 10.92 and mean cut-off at 7.77 was used to divide the tumors into AR high and AR low. 998/1904 (52%) were AR high and as observed in our cohort, these tumors had favourable clinicopathological feature like low grade, low stage, smaller T-size and were mostly lymphnode negative and post-menopausal (p<0.05) (Supplementary Table S4). The ER score and proliferation score was calculated by taking the average of the epithelial makers and proliferation related markers respectively. As seen in our cohort, the AR high tumors had a significantly higher ER score (p<0.001) and lower proliferation score (p<0.001) across all tumors. Kaplan Meier Survival analysis showed the tumors with high AR driven gene score had significantly better disease-free survival (mean survival time of 232.13 vs 211.46 months, log rank p=0.003) when compared to the AR low tumors in the METABRIC cohort. This trend of AR high tumors having a better survival than the low tumors was observed within the HR+HER2- (mean survival time of 247.43 vs 206.2 months, log rank p<0.0001), but no significant difference in survival between the AR groups was observed within the HER2+ and TNBC subtype.

Further, a subset analysis within the TNBC subtype showed that 51/299 (17%) of the tumors were AR high and had lower grade, mostly post-menopausal (p<0.05), had significantly high ER score and low proliferation score (p<0.0001), but none of the other clinic-pathological characters were significantly different between AR high and low tumors.

#### Calculation of EMT score in METABRIC

Given the availability of wider transcriptomic data in external datasets, we derived a EMT score using previously validated set of 77 genes (39). This is a pan-cancer EMT gene signature derived from 1,934 tumors including breast, lung, colon, ovarian, and bladder cancers (total of 11 cancer types). Briefly, the EMT score is determined by subtracting the mean expression of epithelial markers from the mean expression of mesenchymal markers for each sample. A more mesenchymal expression pattern is associated with higher EMT scores. EMT score for was derived for all tumors and compared between the AR high and low tumors. It was observed that the tumors with high AR gene score had significantly higher EMT score (Figure 6A and 6B). This trend was seen in all tumors(p<2.2e-16) as well as in TNBC subtype(p=2.6e-06).

**FIGURE 6:**
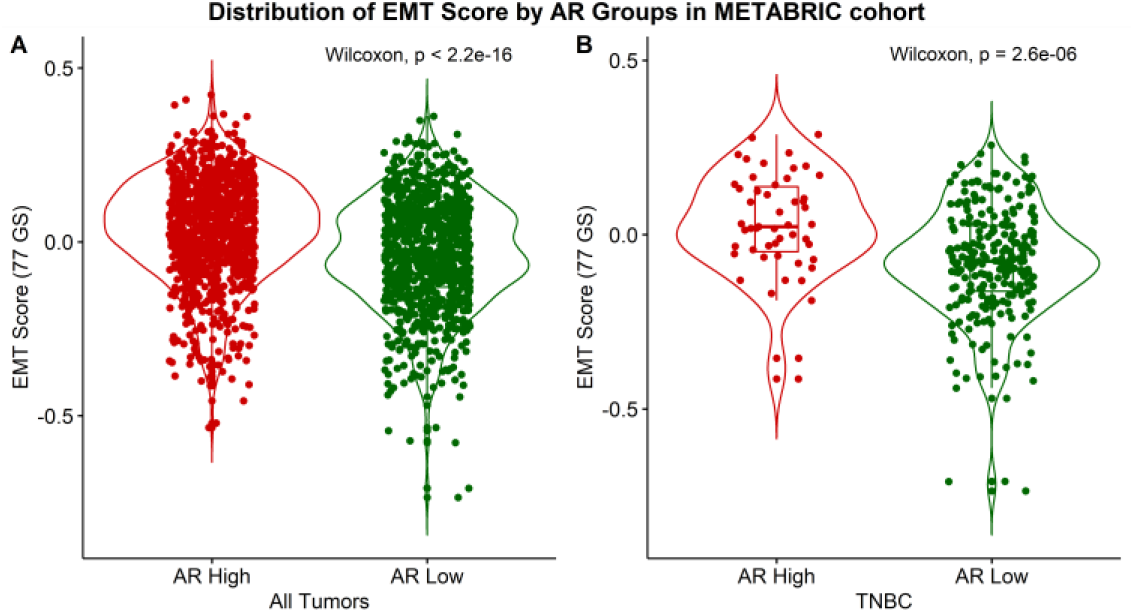
EMT score in AR high and low groups in METABRIC cohort. (A) Distribution of EMT score in all tumors (B) Distribution of EMT score in TNBC tumors.

These results further confirmed the findings from our cohort in that TNBC tumors driven by AR show EMT phenotype. In addition, levels of enzyme 5α Reductase (SRD5A1) were also significantly higher in the AR high tumors(p=0.001) indicating it is likely to be driven by AR. Similar trends were also observed in TCGA cohort (details in the Supplementary data S5).

## Discussion

The role of AR in breast cancer is very complex, context dependent based on ER status and studies have shown its dual behaviour as a promoter of tumor growth in TNBC and inhibitor of tumor progression in HR+HER2-tumors(3,40,41). The prognostic significance of AR and its protein status in TNBC has been reported in a recent multi-institutional study(11). Results suggest AR status alone is not a reliable marker, expression is population specific and its role as a prognostic indicator is highly variable across different cohorts, indicating the need to derive AR driven signatures for better identification of TNBC tumors which could respond to anti-AR therapies. Expression of AR being observed in less than half of the tumors of BC from Indian cohorts(42,43), we attempted to derive an AR driven score using expression profiles of AR regulated genes. A systematic method to create bioinformatic pipeline was used to arrive at the AR regulated genes using publicly available AR+ cell lines representing ER positive (ZR-75-1) and ER negative groups (MDA-MB-453), treated with the non-aromatizable androgen, DHT. This approach is unique as it takes into consideration the presence of ER which is known to highly influence the functional consequence of AR mediated signalling due to their crosstalk(44). Public data sets of cell lines treated under controlled settings were used to simulate physiological conditions. Use of gene expression profiles driven by AR have been largely confined to identification of luminal androgen receptor(LAR) subtype of TNBC and molecular apocrine tumors(45–48) within BC. Like in previous studies, our method derived extensive gene sets, was narrowed down to smaller set of markers to achieve the advantage of easier application in clinical settings. In our cohort, the AR driven gene score identified 36% of the TNBC tumors to have a high AR score, whereas in METABRIC and TCGA, only 17% of the TNBC tumors had high AR score indicating population differences in the molecular composition of TNBC based on ethnicity.

Expression of AR has been associated with better survival in both DFS and overall survival in large datasets within ER positive tumors(49). Similar results were obtained with AR driven high score tumors, showing better survival, favourable clinic-pathological and luminal features with lower proliferation score. Similar results with ER driven features and low proliferation were observed in TNBC tumors.

EMT which is a developmentally conserved complex process, plays a central role in tumor progression, aggression, invasion, metastasis and resistance to therapy. Role of AR and androgens in inducing EMT has been well established in experimental systems of prostate cancer (50,51). Though the regulatory role of AR in EMT was initially disputed, more recent studies have alluded to involving transcription factors such as Slug(52) its regulation of ELF5A2(53,54) involvement of splice variants of AR in inducing EMT and stemness in prostate cancer(55).

Evidence to the induction of EMT by steroid nuclear receptors in BC was referred to by *in silico* approaches(56) and a more definitive mechanism was shown in experimental model system involving MDA-MB 453, a cell line representing the luminal androgen receptor subtype of TNBC. These cells acquired mesenchymal features when treated with AR agonist(13), mediated through β catenin and Wnt signaling and more recent work by the same group show regulation of EMT by RGS2, an AR mediated protein(15). Another group showed overexpression of AR in MDAMB 231 cells, could induce invasiveness through AR/SRC/PI3K complex(57). Graham et al. showed that ZEB1 and AR regulate each other to promote cell migration or EMT in TNBC cell lines (MDA-MB-231 and MDA-MB-453), and a suppressive effect of anti-AR drug bicalutamide on ZEB1 expression(38). Our results from cohort of primary BC showed AR driven tumors had a higher expression of EMT markers within the TNBC subtype and validation in larger public datasets, further confirm the association of AR with this process. Though the positive association between AR and EMT was observed across all tumors, subset analysis showed the phenomenon was confined to TNBC tumors alone. The number of AR driven tumors was higher in HR+HER- and HER2+ tumors, but this trend was not observed in these subtypes. Ahram et al observed treatment with AR agonist DHT on MDA-MB-453(14) induced only partial EMT with changes in morphology from epithelial to mesenchymal phenotype but less changes in migration. Association of AR high group of tumors with better survival despite being associated with higher EMT features, perhaps is due to inability of AR in aiding invasion and migration which are crucial steps in metastases.

ZEB1, major transcription factor known to drive EMT has been shown to have crosstalk and reciprocal relationship with AR in TNBC cells(38). Though we observed trends of higher expression of ZEB1 within AR protein positive tumors, it did not meet the statistical significance perhaps due to lower number of AR positive tumors within the breast cores examined. We could not derive AR driven score in these tumors due to the lack of availability of gene expression in this cohort.

Role of AR in promoting metastasis is quite complex though studies have shown that AR is the only sex steroid receptor detectable in BC metastases and consistent AR expression was noted between primary tumors and its corresponding distant or local metastatic tumor(58,59). Levels of androgen metabolizing enzyme, SRD5A1 decreased in the lymph node metastasis indicating that mere presence of AR protein may not be associated with its ability to promote the process of EMT and the importance of intracrinal levels of androgens in driving the process. Our results showed the expression of SRD5A1 is significantly higher in these AR driven tumors with high EMT score, indirectly implying that active metabolites of androgens might play a role in inducing EMT in these tumors. Studies in other cancers have shown the involvement of SRD5A1 in cell migration(60), further supporting our finding. As seen in the case of AR driven tumors having a higher EMT score confined only to the TNBC tumors, high SRD5A1 levels was observed only in TNBC and not in the other subtypes.

We have derived the association of AR with EMT features by correlative analysis from cohort of breast cancer patients and lack of explanation on the mechanism by which AR induces EMT within TNBC may be construed as limitation of the study. However, the exact mechanism to which AR induces EMT is still not discernible despite being attempted in various experimental models. Though we derived our observation from a small number of TNBC tumors within our cohort, replication of the results in larger external datasets confirms the validity of the score derived in our study.

## Conclusion

The value of AR in BC as prognostic and predictive marker is elusive mostly due to heterogeneity of the disease and difference in methodologies for detection of AR. Approaches involving multiple downstream markers are better and are likely to identify tumors truly driven by AR than AR protein alone. Our results confirm the dual role of AR in different subtypes of BC and warrant in depth understanding of influence of AR on EMT pathways to advance a more informed targeted approach for successful anti metastatic treatment, specifically in triple negative tumors.

## Reference

1. Siegel RL, Miller KD, Jemal A. Cancer statistics, 2018. CA; A Cancer Journal for Clinicians (2018) 68:7–30. doi: 10.3322/CAAC.21442

2. Kittaneh M, Montero AJ, Glück S. Molecular Profiling for Breast Cancer: A Comprehensive Review. Biomarkers in Cancer (2013) 5:61. doi: 10.4137/BIC.S9455

3. Hickey TE, Selth LA, Chia KM, Laven-Law G, Milioli HH, Roden D, Jindal S, Hui M, Finlay-Schultz J, Ebrahimie E, et al. The androgen receptor is a tumor suppressor in estrogen receptor-positive breast cancer. Nat Med (2021) 27:310–320. doi: 10.1038/S41591-020-01168-7

4. Aleskandarany MA, Abduljabbar R, Ashankyty I, Elmouna A, Jerjees D, Ali S, Buluwela L, Diez-Rodriguez M, Caldas C, Green AR, et al. Prognostic significance of androgen receptor expression in invasive breast cancer: transcriptomic and protein expression analysis. Breast Cancer Res Treat (2016) 159:215–227. doi: 10.1007/S10549-016-3934-5

5. Liu YX, Zhang KJ, Tang LL. Clinical significance of androgen receptor expression in triple negative breast cancer-an immunohistochemistry study. Oncology Letters (2018) 15:10008. doi: 10.3892/OL.2018.8548

6. Hwang KT, Kim YA, Kim J, Park JH, Choi IS, Hwang KR, Chai YJ, Park JH. Influence of Androgen Receptor on the Prognosis of Breast Cancer. Journal of Clinical Medicine (2020) 9: doi: 10.3390/JCM9041083

7. Elebro K, Bendahl PO, Jernström H, Borgquist S. Androgen receptor expression and breast cancer mortality in a population-based prospective cohort. Breast Cancer Res Treat (2017) 165:645–657. doi: 10.1007/S10549-017-4343-0

8. Kraby MR, Valla M, Opdahl S, Haugen OA, Sawicka JE, Engstrøm MJ, Bofin AM. The prognostic value of androgen receptors in breast cancer subtypes. Breast Cancer Res Treat (2018) 172:283–296. doi: 10.1007/S10549-018-4904-X

9. Kumar V, Yu J, Phan V, Tudor IC, Peterson A, Uppal H. Androgen Receptor Immunohistochemistry as a Companion Diagnostic Approach to Predict Clinical Response to Enzalutamide in Triple-Negative Breast Cancer. https://doi.org/101200/PO1700075 (2017) 1–19. doi: 10.1200/PO.17.00075

10. Ricciardelli C, Bianco-Miotto T, Jindal S, Butler LM, Leung S, McNeil CM, O’Toole SA, Ebrahimie E, Millar EKA, Sakko AJ, et al. The Magnitude of Androgen Receptor Positivity in Breast Cancer Is Critical for Reliable Prediction of Disease Outcome. Clin Cancer Res (2018) 24:2328–2341. doi: 10.1158/1078-0432.CCR-17-1199

11. Bhattarai S, Klimov S, Mittal K, Krishnamurti U, Li XB, Oprea-Ilies G, Wetherilt CS, Riaz A, Aleskandarany MA, Green AR, et al. Prognostic Role of Androgen Receptor in Triple Negative Breast Cancer: A Multi-Institutional Study. Cancers (Basel) (2019) 11: doi: 10.3390/CANCERS11070995

12. Zhu M-L, Kyprianou N. Role of androgens and the androgen receptor in epithelial-mesenchymal transition and invasion of prostate cancer cells. The FASEB Journal (2010) 24:769–777. doi: 10.1096/FJ.09-136994

13. Ahram M, Bawadi R, Abdullah MS, Alsafadi DB, Abaza H, Abdallah S, Mustafa E. Involvement of β-catenin in Androgen-induced Mesenchymal Transition of Breast MDA-MB-453 Cancer Cells. https://doi.org/101080/0743580020211895829 (2021) 46:114–128. doi: 10.1080/07435800.2021.1895829

14. Ahram M, Abdullah MS, Mustafa SA, Alsafadi DB, Battah AH. Androgen downregulates desmocollin-2 in association with induction of mesenchymal transition of breast MDA-MB-453 cancer cells. Cytoskeleton (2021) 78:391–399. doi: 10.1002/CM.21691

15. Alsafadi DB, Abdullah MS, Bawadi R, Ahram M. The Association of RGS2 and Slug in the Androgen-induced Acquisition of Mesenchymal Features of Breast MDA-MB-453 Cancer Cells. Endocr Res (2022)1-16. doi: 10.1080/07435800.2022.2036752

16. CYP17 Lyase and Androgen Receptor Inhibitor Treatment With Seviteronel Trial (INO-VT-464-006; NCT02580448) - No Study Results Posted - ClinicalTrials.gov. https://clinicaltrials.gov/ct2/show/results/NCT02580448 [Accessed November 1, 2020]

17. Efficacy and Safety of GTx-024 in Patients With ER+/AR+ Breast Cancer - Full Text View - ClinicalTrials.gov. https://clinicaltrials.gov/ct2/show/NCT02463032 [Accessed November 1, 2020]

18. Overmoyer B, Sanz-Altamira P, Taylor RP, Hancock ML, Dalton JT, Johnston MA, Steiner MS. Enobosarm: A targeted therapy for metastatic, androgen receptor positive, breast cancer. Journal of Clinical Oncology (2014) 32:568–568. doi: 10.1200/jco.2014.32.15_suppl.568

19. Arce-Salinas C, Riesco-Martinez MC, Hanna W, Bedard P, Warner E. Complete response of metastatic androgen receptor-positive breast cancer to bicalutamide: Case report and review of the literature. Journal of Clinical Oncology (2016) 34:e21–e24. doi: 10.1200/JCO.2013.49.8899

20. Schaafsma E, Zhang B, Schaafsma M, Tong CY, Zhang L, Cheng C. Impact of Oncotype DX testing on ER+ breast cancer treatment and survival in the first decade of use. Breast Cancer Research (2021) 23:1–11. doi: 10.1186/S13058-021-01453-4/TABLES/2

21. Soliman H, Shah V, Srkalovic G, Mahtani R, Levine E, Mavromatis B, Srinivasiah J, Kassar M, Gabordi R, Qamar R, et al. MammaPrint guides treatment decisions in breast Cancer: Results of the IMPACt trial. BMC Cancer (2020) 20:1–13. doi: 10.1186/S12885-020-6534-Z/TABLES/4

22. Jensen MB, Lænkholm AV, Nielsen TO, Eriksen JO, Wehn P, Hood T, Ram N, Buckingham W, Ferree S, Ejlertsen B. The Prosigna gene expression assay and responsiveness to adjuvant cyclophosphamide-based chemotherapy in premenopausal high-risk patients with breast cancer. Breast Cancer Research (2018) 20:1–10. doi: 10.1186/S13058-018-1012-0/FIGURES/4

23. Need EF, Selth LA, Trotta AP, Leach DA, Giorgio L, O’Loughlin MA, Smith E, Gill PG, Ingman W v., Graham JD, et al. The unique transcriptional response produced by concurrent estrogen and progesterone treatment in breast cancer cells results in upregulation of growth factor pathways and switching from a Luminal A to a Basal-like subtype. BMC Cancer (2015) 15: doi: 10.1186/S12885-015-1819-3

24. Ni M, Chen Y, Lim E, Wimberly H, Bailey ST, Imai Y, Rimm DL, Shirley Liu X, Brown M. Targeting androgen receptor in estrogen receptor-negative breast cancer. Cancer Cell (2011) 20:119–131. doi: 10.1016/J.CCR.2011.05.026

25. Ritchie ME, Phipson B, Wu D, Hu Y, Law CW, Shi W, Smyth GK. limma powers differential expression analyses for RNA-sequencing and microarray studies. Nucleic Acids Research (2015) 43:e47–e47. doi: 10.1093/NAR/GKV007

26. Irizarry RA, Hobbs B, Collin F, Beazer-Barclay YD, Antonellis KJ, Scherf U, Speed TP. Exploration, normalization, and summaries of high density oligonucleotide array probe level data. Biostatistics (2003) 4:249–264. doi: 10.1093/BIOSTATISTICS/4.2.249

27. Schaefer MH, Fontaine JF, Vinayagam A, Porras P, Wanker EE, Andrade-Navarro MA. HIPPIE: Integrating protein interaction networks with experiment based quality scores. PLoS One (2012) 7: doi: 10.1371/JOURNAL.PONE.0031826

28. Alanis-Lobato G, Andrade-Navarro MA, Schaefer MH. HIPPIE v2.0: enhancing meaningfulness and reliability of protein-protein interaction networks. Nucleic Acids Research (2017) 45:D408. doi: 10.1093/NAR/GKW985

29. Shannon P, Markiel A, Ozier O, Baliga NS, Wang JT, Ramage D, Amin N, Schwikowski B, Ideker T. Cytoscape: a software environment for integrated models of biomolecular interaction networks. Genome Res (2003) 13:2498–2504. doi: 10.1101/GR.1239303

30. Rakshit H, Rathi N, Roy D. Construction and analysis of the protein-protein interaction networks based on gene expression profiles of Parkinson’s disease. PLoS One (2014) 9: doi: 10.1371/JOURNAL.PONE.0103047

31. Yu G, Li F, Qin Y, Bo X, Wu Y, Wang S. GOSemSim: an R package for measuring semantic similarity among GO terms and gene products. Bioinformatics (2010) 26:976–978. doi: 10.1093/BIOINFORMATICS/BTQ064

32. Sankala H, Vaughan C, Wang J, Deb S, Graves PR. Upregulation of the mitochondrial transport protein, Tim50, by mutant p53 contributes to cell growth and chemoresistance. Arch Biochem Biophys (2011) 512:52–60. doi: 10.1016/J.ABB.2011.05.005

33. Rajarajan S, Korlimarla A, Alexander A, Anupama CE, Ramesh R, Srinath BS, Sridhar TS, Prabhu JS. Pre-Menopausal Women With Breast Cancers Having High AR/ER Ratios in the Context of Higher Circulating Testosterone Tend to Have Poorer Outcomes. Frontiers in Endocrinology (2021) 12:1. doi: 10.3389/FENDO.2021.679756/FULL

34. Korlimarla A, Prabhu JS, Anupama CE, Remacle J, Wahi K, Sridhar TS. Separate qualitycontrol measures are necessary for estimation of RNA and methylated DNA from formalin-fixed, paraffin-embedded specimens by quantitative PCR. J Mol Diagn (2014) 16:253–260. doi: 10.1016/J.JMOLDX.2013.11.003

35. Prabhu JS, Korlimarla A, Anupama CE, Alexander A, Raghavan R, Kaul R, Desai K, Rajarajan S, Manjunath S, Correa M, et al. Dissecting the Biological Heterogeneity within Hormone Receptor Positive HER2 Negative Breast Cancer by Gene Expression Markers Identifies Indolent Tumors within Late Stage Disease. Transl Oncol (2017) 10:699–706. doi: 10.1016/J.TRANON.2017.04.011

36. Prabhu JS, Korlimarla A, Desai K, Alexander A, Raghavan R, Anupama CE, Dendukuri N, Manjunath S, Correa M, Raman N, et al. A Majority of Low (1-10%) ER Positive Breast Cancers Behave Like Hormone Receptor Negative Tumors. J Cancer (2014) 5:156–165. doi: 10.7150/JCA.7668

37. J G, BA A, U D, G D, B G, SO S, Y S, A J, R S, E L, et al. Integrative analysis of complex cancer genomics and clinical profiles using the cBioPortal. Sci Signal (2013) 6: doi: 10.1126/SCISIGNAL.2004088

38. Graham TR, Yacoub R, Taliaferro-Smith L, Osunkoya AO, Odero-Marah VA, Liu T, Kimbro KS, Sharma D, O’Regan RM. Reciprocal regulation of ZEB1 and AR in triple negative breast cancer cells. Breast Cancer Res Treat (2010) 123:139–147. doi: 10.1007/S10549-009-0623-7

39. Mak MP, Tong P, Diao L, Cardnell RJ, Gibbons DL, William WN, Skoulidis F, Parra ER, Rodriguez-Canales J, Wistuba II, et al. A Patient-Derived, Pan-Cancer EMT Signature Identifies Global Molecular Alterations and Immune Target Enrichment Following Epithelial-to-Mesenchymal Transition. Clin Cancer Res (2016) 22:609–620. doi: 10.1158/1078-0432.CCR-15-0876

40. Choi JE, Kang SH, Lee SJ, Bae YK. Androgen Receptor Expression Predicts Decreased Survival in Early Stage Triple-Negative Breast Cancer. Annals of Surgical Oncology 2014 22:1 (2014) 22:82–89. doi: 10.1245/S10434-014-3984-Z

41. Asano Y, Kashiwagi S, Onoda N, Kurata K, Morisaki T, Noda S, Takashima T, Ohsawa M, Kitagawa S, Hirakawa K. Clinical verification of sensitivity to preoperative chemotherapy in cases of androgen receptor-expressing positive breast cancer. British Journal of Cancer 2016 114:1 (2016) 114:14–20. doi: 10.1038/bjc.2015.434

42. Vellaisamy G, Tirumalae R, Inchara Y. Expression of androgen receptor in primary breast carcinoma and its relation with clinicopathologic features, estrogen, progesterone, and her-2 receptor status. Journal of Cancer Research and Therapeutics (2019) 15:989–993. doi: 10.4103/jcrt.JCRT_572_17

43. Anand A, Singh KR, Kumar S, Husain N, Kushwaha JK, Sonkar AA. Androgen Receptor Expression in an Indian Breast Cancer Cohort with Relation to Molecular Subtypes and Response to Neoadjuvant Chemotherapy - A Prospective Clinical Study. Breast Care (2017) 12:160–164. doi: 10.1159/000458433

44. Need EF, Selth LA, Harris TJ, Birrell SN, Tilley WD, Buchanan G. Research resource: interplay between the genomic and transcriptional networks of androgen receptor and estrogen receptor α in luminal breast cancer cells. Mol Endocrinol (2012) 26:1941–1952. doi: 10.1210/ME.2011-1314

45. Farmer P, Bonnefoi H, Becette V, Tubiana-Hulin M, Fumoleau P, Larsimont D, MacGrogan G, Bergh J, Cameron D, Goldstein D, et al. Identification of molecular apocrine breast tumours by microarray analysis. Oncogene (2005) 24:4660–4671. doi: 10.1038/SJ.ONC.1208561

46. Alvarez J, Schaffer M, Karkera J, Martinez G, Gaffney D, Bell K, Sharp M, Wong J, Hertzog B, Ricci D, et al. Abstract P5-01-09: Identification of Molecular Apocrine Triple Negative Breast Cancer Using a Novel 2-Gene Assay and Comparison with Androgen Receptor Protein Expression and Gene Expression Profiling by DASL. Cancer Research (2012) 72:P5-01-09. doi: 10.1158/0008-5472.SABCS12-P5-01-09

47. Rody A, Karn T, Liedtke C, Pusztai L, Ruckhaeberle E, Hanker L, Gaetje R, Solbach C, Ahr A, Metzler D, et al. A clinically relevant gene signature in triple negative and basal-like breast cancer. Breast Cancer Research (2011) 13:1–12. doi: 10.1186/BCR3035/TABLES/4

48. Burstein MD, Tsimelzon A, Poage GM, Covington KR, Contreras A, Fuqua SAW, Savage MI, Osborne CK, Hilsenbeck SG, Chang JC, et al. Comprehensive genomic analysis identifies novel subtypes and targets of triple-negative breast cancer. Clin Cancer Res (2015) 21:1688–1698. doi: 10.1158/1078-0432.CCR-14-0432

49. Kensler KH, Poole EM, Heng YJ, Collins LC, Glass B, Beck AH, Hazra A, Rosner BA, Eliassen AH, Hankinson SE, et al. Androgen Receptor Expression and Breast Cancer Survival: Results From the Nurses’ Health Studies. JNCI Journal of the National Cancer Institute (2019) 111:700. doi: 10.1093/JNCI/DJY173

50. Zhu M-L, Kyprianou N. Role of androgens and the androgen receptor in epithelial-mesenchymal transition and invasion of prostate cancer cells. The FASEB Journal (2010) 24:769–777. doi: 10.1096/FJ.09-136994

51. Matuszak EA, Kyprianou N. Androgen regulation of epithelial-mesenchymal transition in prostate tumorigenesis. Expert Rev Endocrinol Metab (2011) 6:469–482. doi: 10.1586/EEM.11.32

52. Wu K, Gore C, Yang L, Fazli L, Gleave M, Pong RC, Xiao G, Zhang L, Yun EJ, Tseng SF, et al. Slug, a unique Androgen-regulated transcription factor, coordinates androgen receptor to facilitate castration resistance in prostate cancer. Molecular Endocrinology (2012) 26:1496–1507. doi: 10.1210/ME.2011-1360

53. Zheng Y, Li P, Huang H, Ye X, Chen W, Xu G, Zhang F. Androgen receptor regulates eIF5A2 expression and promotes prostate cancer metastasis via EMT. Cell Death Discovery 2021 7:1 (2021) 7:1–8. doi: 10.1038/s41420-021-00764-x

54. Papanikolaou S, Vourda A, Syggelos S, Gyftopoulos K. Cell Plasticity and Prostate Cancer: The Role of Epithelial-Mesenchymal Transition in Tumor Progression, Invasion, Metastasis and Cancer Therapy Resistance. Cancers (Basel) (2021) 13: doi: 10.3390/CANCERS13112795

55. Ojo D, Lin X, Wong N, Gu Y, Tang D. Prostate Cancer Stem-like Cells Contribute to the Development of Castration-Resistant Prostate Cancer. Cancers (Basel) (2015) 7:2290. doi: 10.3390/CANCERS7040890

56. Voutsadakis IA. Epithelial-Mesenchymal Transition (EMT) and Regulation of EMT Factors by Steroid Nuclear Receptors in Breast Cancer: A Review and in Silico Investigation. Journal of Clinical Medicine (2016) 5: doi: 10.3390/JCM5010011

57. Giovannelli P, di Donato M, Auricchio F, Castoria G, Migliaccio A. Androgens Induce Invasiveness of Triple Negative Breast Cancer Cells Through AR/Src/PI3-K Complex Assembly. Scientific Reports (2019) 9: doi: 10.1038/s41598-019-41016-4

58. Shibahara Y, Miki Y, Sakurada C, Uchida K, Hata S, McNamara K, Yoda T, Takagi K, Nakamura Y, Suzuki T, et al. Androgen and androgen-metabolizing enzymes in metastasized lymph nodes of breast cancer. Human Pathology (2013) 44:2338–2345. doi: 10.1016/J.HUMPATH.2013.04.021

59. McNamara KM, Yoda T, Miki Y, Nakamura Y, Suzuki T, Nemoto N, Miyashita M, Nishimura R, Arima N, Tamaki K, et al. Androgen receptor and enzymes in lymph node metastasis and cancer reoccurrence in triple-negative breast cancer. Int J Biol Markers (2015) 30:e184–e189. doi: 10.5301/JBM.5000132

60. Wei R, Zhong S, Qiao L, Guo M, Shao M, Wang S, Jiang B, Yang Y, Gu C. Steroid 5α-Reductase Type I Induces Cell Viability and Migration via Nuclear Factor-κB/Vascular Endothelial Growth Factor Signaling Pathway in Colorectal Cancer. Frontiers in Oncology (2020) 10:1501. doi: 10.3389/FONC.2020.01501

