## Supplementary data and tables for "Androgen Receptor driven gene score identifies tumors with Epithelial to Mesenchymal Transition features in triple negative breast cancer"

**Supplementary Data S1: Hormonal treatment with dose details and duration of treatment is available in excel sheet S1.**

**Supplementary Table S2: Clinicopathological features of all patients in our cohort (N=244)**

| **Clinicopathological**  **characteristics** |  | **All samples**  **(N=244)** |
| --- | --- | --- |
|  |  | N (%) |
| Age | Mean | 56 |
|  | Median | 56 |
| T size | Mean | 3.34 |
|  | Median | 3 |
|  | 1 | 66(28) |
|  | 2 | 142(60) |
|  | 3 | 27(12) |
| Lymph Node | Positive | 144(61) |
|  | Negative | 92(39) |
| Stage | I | 36(15) |
|  | II | 120(49) |
|  | III | 78(32) |
|  | IV | 10(4) |
| Grade | I | 18(10) |
|  | II | 111(55) |
|  | III | 102(35) |
| Estrogen Receptor | Positive | 170(70) |
|  | Negative | 74(30) |
| Progesterone Receptor | Positive | 158(65) |
|  | Negative | 86(35) |
| HER2 | Positive | 51(21) |
|  | Negative | 167(68) |
|  | Equivocal | 26(8) |

Menopausal status	Pre	69(28)

Post	175(72)

**Supplementary Table S3**: Details of primer sequences

| **Gene** | **Primer Sequence** | **Product Size(bp)** | **NM_sequence**  **ID** | **Gene Details** |
| --- | --- | --- | --- | --- |
| CYP4Z1 | F- gctggattaaggaactgcattgggc | 88 | NM_178134.3 | Cytochrome P450 Family 4 Subfamily Z Member |
|  | R- ccagcttgaagcggagcagagtta |  |  |  |
| ABCC11 | F- tcaacggaaatgctgtgccgga | 85 | NM_001370496.1 | ATP Binding Cassette Subfamily C Member 11 |
|  | R- agtggtggtagaccctccaactca |  |  |  |
| TFAP2B | F- gagtgacctgcactcccgaaagaa | 83 | NM_003221.4 | Transcription Factor AP-2 Beta |
|  | R-ggtcctgcgccagtagatccgtaaat |  |  |  |
| SOCS2 | F-catcccttctctctctgccaccattt | 80 | NM_001270467.2 | Suppressor Of Cytokine Signaling 2 |
|  | R-cgtccttccttgaagtcagtgcgaat |  |  |  |
| GADD45G | F-tggagaagctcagcctgttt | 109 | NM_006705.4 | Growth Arrest And DNA Damage Inducible Gamma |
|  | R-acgtcgatcagaccaaggtc |  |  |  |
| ZNF689 | F-catcttccggtgcctttcac | 100 | NM_138447.3 | Zinc Finger Protein 689 |
|  | R-cggagggaatcggaattggt |  |  |  |
| ID1 | F-tgcttcgggcttccacctcatttt | 85 | NM_002165.4 | Inhibitor Of DNA Binding 1, HLH Protein |
|  | R-ctgccactggcgactttcatgattct |  |  |  |
| PIP | F- ctctgcctgcagttggggg | 100 | NM_002652.3 | Prolactin Induced Protein |
|  | R- cttcgtcatttggacgtactgacttgg |  |  |  |
| UGT2B11 | F-gctgctggctgcgctacttaacat | 95 | NM_001073.3 | UDP Glucuronosyltransferase Family 2 Member B11 |
|  | R-ggaaaatcagtcctccactgtgcct |  |  |  |
| SEC14L2 | F-accatgactgaccctgatgg | 96 | NM_001204204.3 | SEC14 Like Lipid Binding 2 |
|  | R-tttcacctggtctcgcacat |  |  |  |
| DOCK2 | F-acatcccatctgtcctgcat | 91 | NM_004946.3 | Dedicator Of Cytokinesis 2 |
|  | R-ggagggatgcaggtgtagaa |  |  |  |
| KCNMA1 | F-actgcagccgggttcatccatt | 89 | NM_001014797.3 | Potassium Calcium-Activated Channel Subfamily M Alpha 1 |
|  | R-ttcccagtaggtgagagagcctggtt |  |  |  |
| CYP19A1 | F-acgctcttcttgaggatccc | 89 | NM_000103.4 | Cytochrome P450 Family 19 Subfamily A Member 1 |
|  | R-ttgatgaggagagcttgcca |  |  |  |
| SRD5A1 | F-cttgagccattgtgcagtgt | 88 | NM_001047.4 | Steroid 5 Alpha-Reductase 1 |
|  | R-caacatgcccgttaaccaca |  |  |  |
| UBE2C | F- tgccctgtatgatgtcagga | 73 | NM_001281741.2 | Ubiquitin Conjugating Enzyme E2 C |
|  | R- gggactatcaatgttgggttctc |  |  |  |
| ANLN | F-acagccactttcagaagcaag | 73 | NM_001284301.3 | Anillin Actin Binding Protein |
|  | R-cgatggttttgtacaagatttctc |  |  |  |
| CENPF | F-gtggcagcagatcacaa | 86 | NM_016343.4 | Centromere Protein F |
|  | R-ggatttcgtggtgggttc |  |  |  |
| BIRC5 | F-gcacaaagccattctaagtc | 82 | NM_001012270.2 | Baculoviral IAP Repeat Containing 5 |
|  | R-gacgcttcctatcactctattc |  |  |  |
| TWIST1 | F-ccggagacctagatgtcattg | 109 | NM_000474.4 | Twist Family BHLH Transcription Factor 1 |
|  | R-tttccaagaaaatctttggcata |  |  |  |
| SNAI2 | F-ccaaactacagcgaactgga | 82 | NM_003068.5 | Snail Family Transcriptional Repressor 2 |
|  | R-gtggtatgacaggcatggag |  |  |  |
| ZEB1 | F-gcacctgaagaggaccagag | 72 | NM_001128128.3 | Zinc Finger E-Box Binding Homeobox 1 |
|  | R-tgcatctggtgttccatttt |  |  |  |
| ZEB2 | F-tgcacagagtgtggcaaggc | 90 | NM_001171653.2 | Zinc Finger E-Box Binding Homeobox 2 |
|  | R-tgggcactcgtaaggtttttcacc |  |  |  |

**Supplementary Table S4: Comparison of clinical variables between high and low AR groups in METABRIC cohort**

| **Clinicopathological**  **characteristics** |  | **AR high**  **(N=998)** | **AR Low**  **(N=906)** | **p-value** |
| --- | --- | --- | --- | --- |
|  |  | N (%) | N (%) |  |
| Age | Mean | 62.41 | 59.62 | **<0.0001*** |
|  | Median | 63.02 | 60.4 |  |
| T size | Mean | 2.54 | 2.7 | **0.022*** |
|  | Median | 2.2 | 2.4 |  |
|  | 1 | 449(45) | 375(42) |  |
|  | 2 | 494(50) | 475(53) |  |
|  | 3 | 45(5) | 48(5) |  |
| Lymph Node | Positive | 451(45) | 460(51) | **0.015*** |
|  | Negative | 547(55) | 446(49) |  |
| Stage | I | 267(36) | 208(31) |  |
|  | II | 404(56) | 396(59) | **0.039*** |
|  | III | 49(7) | 66(10) |  |
|  | IV | 4(1) | 5(1) |  |
| Grade | I | 125(14) | 34(5) | **<0.0001*** |
|  | II | 440(48) | 271(32) |  |
|  | III | 350(38) | 544(64) |  |
| Estrogen Receptor | Positive | 847(85) | 612(68) | **<0.0001*** |
|  | Negative | 151(15) | 294(32) |  |
| Progesterone Receptor | Positive | 617(62) | 392(43) | **<0.0001*** |
|  | Negative | 381(38) | 514(57) |  |
| HER2 | Positive | 156(15) | 80(9) | **<0.0001*** |
|  | Negative | 842(85) | 826(91) |  |

Menopausal status	Pre	186(19)	225(25)	**0.001***

Post	812(81)	681(75)

### **Supplementary Data S5**

We next accessed the TCGA database (https://www.cancer.gov/tcga). to validate our findings. From a total of 1084 tumors, 977 tumors had gene expression values for all the six AR driven genes. Of the 977 tumors, 543 tumors were HR+HER2-, 107 were TNBC and 154 were HER2+ and the rest of the tumors were non-categorizable. The six AR driven genes were significantly and positively correlated with AR transcript and the AR driven gene score ranged from 2.89 to 13.07.A mean cut-off at 8.46 was used to divide the tumors into AR high and AR low. 542/977 (55%) were AR high and these tumors had a higher ER score (p<0.0001), lower proliferation score(p<0.0001), smaller T-size(p=0.03), lower stage(p=0.12), higher age(p=0.001) and mostly post-menopausal(p=0.06). Unlike our cohort, Kaplan Meier Survival analysis did not show any survival difference between the two AR groups in the TCGA cohort in all tumors.

Further, a subset analysis within the TNBC subtype showed that 18/107 (17%) of tumors were AR high and both clinical characters and survival was not significantly different between the AR groups. However, the AR high tumors showed significantly higher ER score (p<0.0001) and lower proliferation score (p<0.0001).

As done in the METABRIC cohort, we then used the 77 genes (Mak MP et al ,2016) to calculate the EMT score. The tumors with high AR gene score had significantly higher EMT score in all tumors(p=0.006). Similar trend was seen in TNBC subtype, though not significant (p=0.11) as shown in the figure below.

Further, as observed in our cohort the SRD5A1 levels were significantly higher in the AR driven tumors in the TCGA cohort(p=0.04).

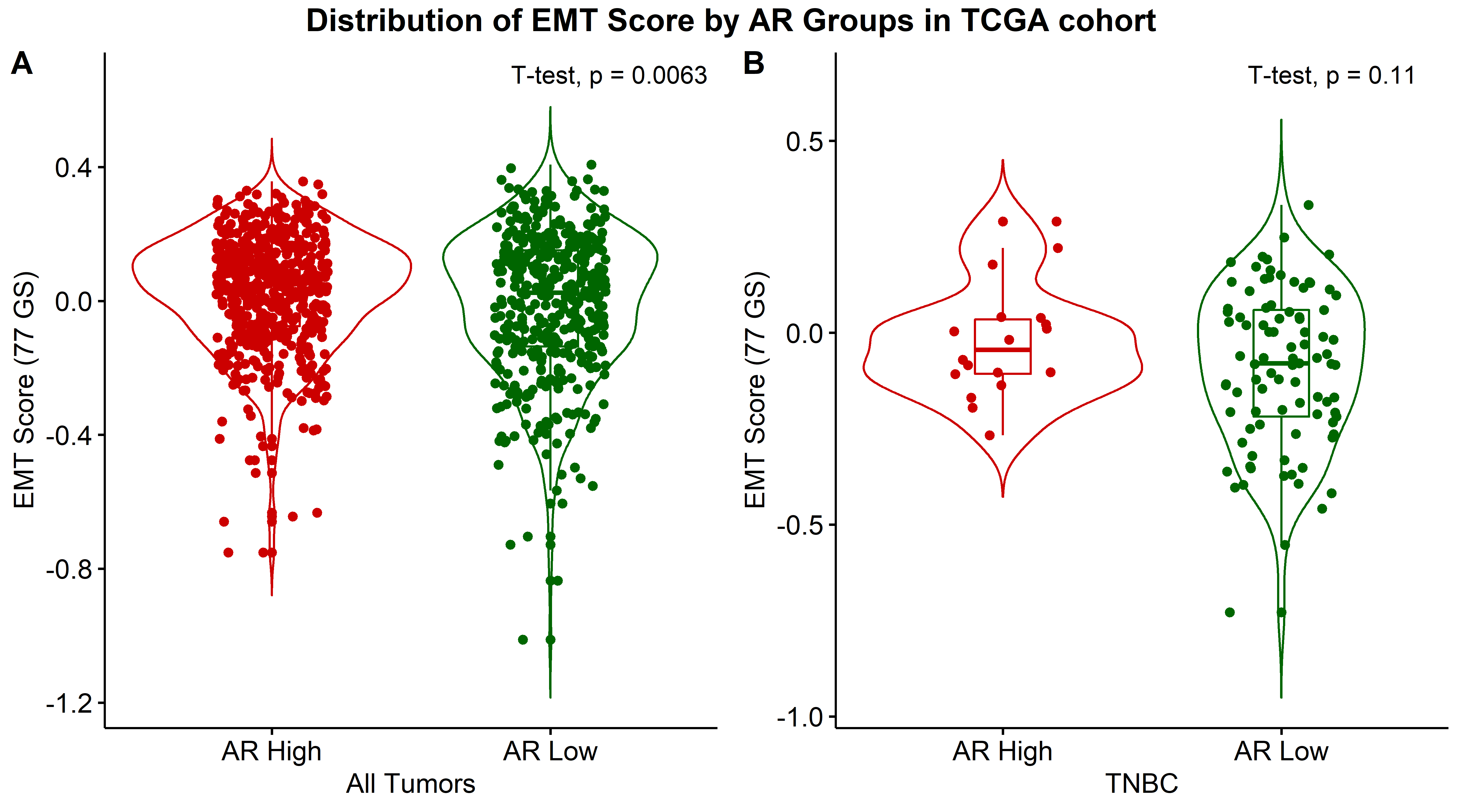

FIGURE S5: EMT score in AR high and low groups in TCGA cohort. (A) Distribution of EMT score in all tumors (B) Distribution of EMT score in TNBC tumors.
